# The *E. coli* DEAD box ATPase CsdA, is a NAD^+^ capped RNA binding protein

**DOI:** 10.64898/2026.06.03.730015

**Authors:** Kaustuv Das, Kathryn G Dzurik, Yogeshwari Singh, Yanbao Yu, Karl R Schmitz, Jared M Schrader, W Seth Childers, Jeremy G Bird

## Abstract

5′ nicotinamide adenine dinucleotide (NAD^+^) caps are one of the most common metabolites derived non-canonical caps reported on bacterial RNAs. Multiple decapping proteins are known to regulate the stability of NAD^+^ capped transcripts. However, no other proteins have been identified that preferentially interact with these NAD^+^ caps, and mechanistic details of the cap-dependent recognition remain poorly understood. Using an affinity capture approach, we identified multiple *E. coli* proteins that selectively recognize NAD^+^ caps, including the ATP-dependent RNA helicase, CsdA. CsdA preferentially interacts directly with NAD^+^ capped RNAs and can discriminate between 5′ NAD^+^ capped and 5′ triphosphate end transcripts. Binding to NAD^+^ capped RNA versus 5′ triphosphate RNA more greatly enhances the ATPase activity of CsdA and the presence of NAD^+^ caps on transcripts modulates the ability of CsdA to form RNA condensates. Furthermore, we find that CsdA enhances the decapping activity of the NADH hydrolase NudC, suggesting CsdA plays a role in regulating the degradation of NAD^+^ capped transcripts. CsdA is the first identified NAD^+^ cap reader protein and its preference for binding NAD^+^ capped RNA provides a mechanism by which *E. coli* cells link RNA stability to the identity of the 5′ cap.

**GRAPHICAL ABSTRACT:** 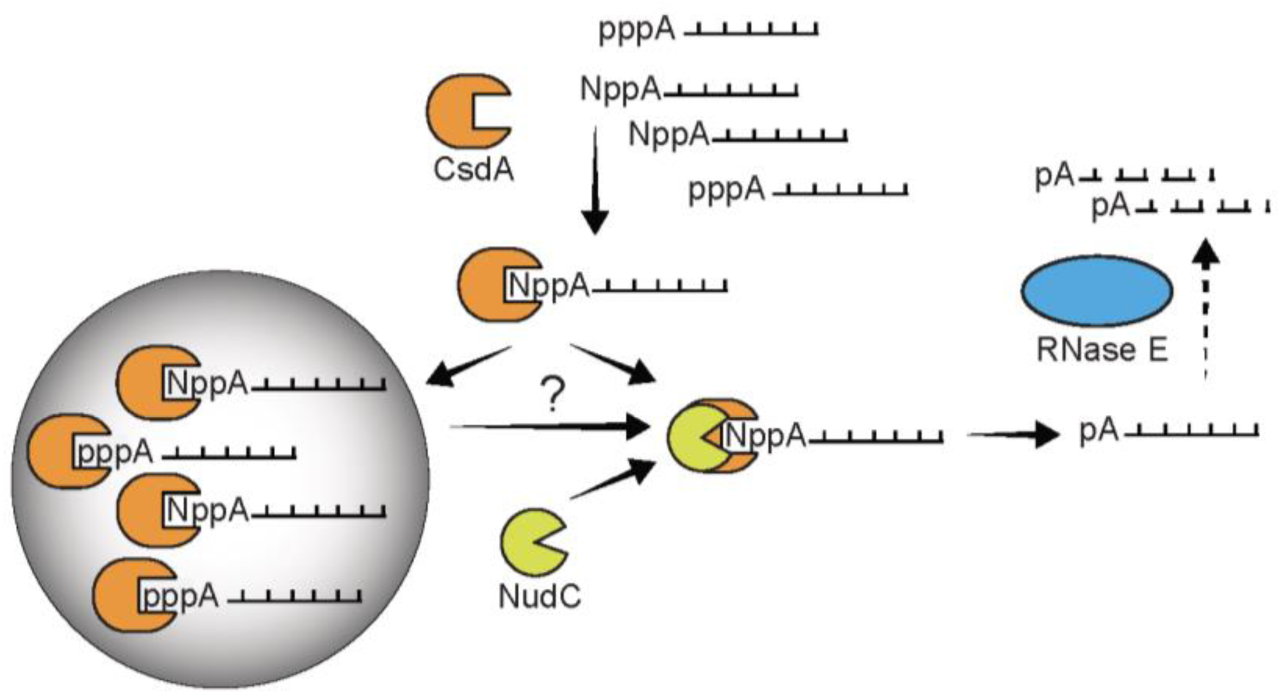

## INTRODUCTION

Nicotinamide adenine dinucleotide (NAD^+^) capped RNA was first identified in bacteria and has been shown to play a role in RNA stabilization in *E. coli* (1,2). Unlike the 7-methylguanylate (m^7^G) cap at 5′ ends of eukaryotic mRNAs, 5′ NAD^+^ caps result from NAD^+^ being incorporated by RNAP as a non-canonical initiating nucleotide (1,2). In *E. coli*, 5′ NAD^+^ caps have been found on both coding and non-coding transcripts (1,3); 16 out of 77 annotated small regulatory RNAs in *E. coli* were shown to be NAD^+^ capped, with levels ranging from 1.6% to 22.4% (4). Interestingly, NAD^+^ capping of the same transcript varies depending on the growth phase of *E. coli* (2,3). NAD^+^ capping of most transcripts was observed to be higher in stationary phase relative to exponential phase (2,3). Importantly, the stability of the NAD^+^ capped fraction of a reporter RNA under the control of the *colE1* origin of replication RNA1 promoter, was found to be enhanced in stationary phase compared to exponential phase (2).

The NAD^+^ cap has been demonstrated to increase the stability of *E. coli* RNAs 2-fold during exponential growth and 4-fold during stationary phase (2). This stability is presumably dependent on protection of the RNA 5′ end by the RNA cap. RNase E, one of the primary components of the RNA degradosome, facilitates 5′ end-dependent RNA turnover in *E. coli* and is unable to degrade NAD^+^ capped RNA *in vitro* (1). This necessitates proteins that can process NAD^+^ caps to generate monophosphate ends, a preferred substrate for RNase E. Both the nudix hydrolase family and non-nudix NAD^+^ decapping enzymes that can cleave or completely remove NAD^+^ from the 5′ end of RNA to generate a monophosphate end have been found across the tree of life. These enzymes include NudC in *E. coli*, NPY1 and DXO in yeast, and Nudt12 in humans (5–7). The existence of these NAD^+^ decapping enzymes indicates an evolutionarily conserved need for the removal of 5′ NAD^+^ caps. NudC, the primary protein that removes RNA 5′ NAD^+^ caps in *E. coli*, is reported to bind RNA in a non-specific manner, and is likely unable to bind selectively to 5′ NAD^+^ capped transcripts *in vivo* (5).

Over the last decade, 5′ NAD^+^ capped RNAs have been reported in organisms from all branches of the tree of life, including archaea, yeast, plants, and human cells suggesting that these metabolite caps may have a universal function (8–11). To date, the only proteins shown to interact with 5′ NAD^+^ caps act directly on the cap itself, either by removing it or, in the case of ModB, a phage ADP-ribosylase, covalently linking 5′ NAD^+^ capped RNA to ribosomal proteins rL2 and rS1 (12,13). The prevalence of NAD^+^ caps across all domains of life supports the existence of a broader NAD^+^ cap interactome, encompassing proteins in addition to canonical decapping enzymes or phage anti-host proteins. In this study, we set out to identify other RNA binding protein(s) that interact specifically with transcripts based on the existence of a 5′ NAD^+^ moiety.

We performed NAD cap RNA affinity purification (NcRAP) (14) with *E. coli* K-12 lysate to pull down and identify RNA binding proteins that specifically interact with NAD^+^ capped RNAs. We identified multiple novel 5′ NAD^+^ cap interactors, including an ATP dependent RNA helicase, CsdA. *E. coli* CsdA plays a key role in ribosome assembly and is essential for growth at lower temperatures, primarily due to its role in RNA degradation (15–17). Consistent with this role, an earlier study reported physical interaction by Co-IP between RNase E and CsdA following a shift of cultures to 15°C (18). More recently, NudC has been shown to physically interact with CsdA at both 37°C and 16°C in the BL21(DE3)-RIL strain of *E. coli* (19).

Here, we demonstrate that CsdA preferentially interacts with NAD^+^ capped RNA over triphosphate end RNA (5′ ppp RNA) with 8-fold higher affinity *in vitro*. Using biochemical assays, we show that CsdA has differential binding kinetics for NAD^+^ capped RNA compared to 5′ ppp RNA. This preferential interaction influences both the ATPase activity and RNA condensate forming activity of CsdA. Functionally, we show that CsdA enhances the efficiency of NAD^+^ decapping by NudC *in vitro*, consistent with previous evidence of a CsdA-NudC interaction (19). This finding supports a model in which CsdA specifically recognizes 5′ NAD^+^ capped transcripts and promotes their decapping by NudC. CsdA is the first identified NAD^+^ cap RNA interacting protein that does not directly act on the 5′ NAD^+^ cap, instead our results support it playing a role in NAD^+^ capped RNA homeostasis in *E. coli*.

## MATERIAL AND METHODS

A complete list of reagents and equipment used in this study can be found in Supplemental Table 1.

### Bacterial Strains

All strains of bacteria used for this study are derivatives of the *E. coli* K-12 MG1655 *strain* and are listed in Supplemental Table 2.

The MG1655 *E. coli csdA-3x FLAG* strain was generated using the scarless genome editing method from Kim *et al.* (2014) (20). Briefly, *E. coli* MG1655 cells transformed with pSLTS plasmid were grown overnight, back diluted 1:100 the following day in LB media and grown for 2 hours before the culture was induced with 2 mM (final concentration) of arabinose to express λ Red recombinase. Post arabinose addition, the cells were grown for a further 2 hours, after which they were harvested and made electrocompetent by washing the cells 4 times with ice cold 10% glycerol as described previously (21). Next, MG1655 pSLTS electrocompetent cells were transformed via electroporation with the dsDNA oligo (KD09, **Supplementary Table 3**), incubated for 3 hours in SOC media (New England Biolabs) at 30°C to facilitate recombination, and plated onto agar plates supplemented with 50 µg/mL kanamycin and 25 µg/mL carbenicillin for selection.

Colonies were re-validated by streaking on plates supplemented with 50 µg/mL kanamycin and 25 µg/mL carbenicillin and incubated overnight at 30°C. To remove the *kanR* cassette, a single colony was resuspended in 500 µL LB media and plated on agar plates supplemented with 25 µg/mL carbenicillin or 25 µg/mL carbenicillin and 100 ng/mL anhydrous tetracycline. The presence of anhydrous tetracycline induced the expression of ISce-I endonuclease from the pSLTS plasmid to make a dsDNA break which was needed to remove the kanamycin cassette (20). Next, roughly 40 colonies from LB plates supplemented with 25 µg/mL carbenicillin and 100 ng/mL anhydrous tetracycline were replica plated onto plain LB plates and LB plates supplemented with 50 µg/mL kanamycin. The cells were incubated at 37°C overnight. Colonies that grew in plain LB but not in LB with kanamycin were the ones that had correctly incorporated 3xFLAG epitope tag incorporated into the csdA coding sequence and were successfully cured of the kanamycin cassette after transformation of pCP20 plasmid, resulting in a scarless addition of a C-terminal 3xFLAG epitope tag on the *csdA* gene. Successful chromosomal editing was confirmed with Bacterial genome sequencing by Oxford Nanopore Technology (Plasmidsaurus) with standard analysis and annotation.

### Plasmid construction

The pET28c-CsdA-His plasmid was constructed by cloning the *csdA* gene sequence (using primers KD07/KD08, **Supplementary Table 3**) between HindIII and Xba1 sites in pET28c plasmid. The pET28c-NudC-His and pET28c-NudCE178Q-His plasmid were generated as described previously (2). The plasmid vector, pSLTS encoding the λ Red recombinase, was obtained from Addgene (plasmid #59386) (20). The pCP20 plasmid was kindly provided by Dr. Bryce Nickels (Rutgers University, New Jersey).

### Purification of CsdA and NudC proteins

*E. coli* CsdA was purified from pET28c-CsdA-His transformed into NiCo21(DE3) cells (New England Biolabs). Transformed cells were grown overnight in 25 mL LB media in presence of 50 µg/mL kanamycin. 1 L of LB media supplemented with 50 µg/mL kanamycin was inoculated with a 1:100 dilution from the overnight cultures and allowed to grow up to an OD ∼ 0.6 at 37°C. Expression of proteins was induced using 0.5 mM IPTG for CsdA and incubated overnight at 16°C. After incubation, cells were pelleted by centrifugation and then resuspended in lysis buffer (50 mM Tris base, 500 mM NaCl, 20 mM imidazole, 0.5 mM EDTA, 1 mM DTT, pH 7.5) followed by sonication and centrifugation at 15000 x g for 30 minutes. The resulting supernatant was incubated with Ni-NTA Agarose beads (MCLAB) for 1 hour at 4°C, after which it was loaded onto a gravity column and washed with 5 column volumes of washing buffer (50 mM Tris base, 500 mM NaCl, 20 mM imidazole, 0.5 mM DTT, pH 7.5). Protein elution from the beads was done using a single volume of 10 mL of elution buffer (50 mM Tris base, 500 mM NaCl, 250 mM imidazole, 0.5 mM DTT, pH 7.5) followed by spin concentration (Amicon Ultra Centrifugal Filter, 30 kDa MWCO). The protein was further purified by size exclusion chromatography (Superdex 200 16/600; Cytiva) in buffer containing 50 mM Tris, 500 mM NaCl, 2 mM DTT, and 5% glycerol. Peak fractions were pooled, spin concentrated and stored at -80°C until use. Final protein purity was determined to be ≥98% pure by SDS-PAGE with Coomassie staining.

*E. coli* Nudix hydrolase protein, NudC and its inactive mutant NudC E178Q, were prepared from pET28c-NudC-His and pET28c-NudCE178Q-His respectively and transformed into NiCo21 (DE3) cells (New England Biolabs) (2). The culture conditions and purification using metal ion affinity chromatography on Ni-NTA agarose, and Superdex S200 16/600 were performed as reported previously (^5^). Final protein purity was determined to be ≥98% pure by SDS-PAGE with Coomassie staining. All protein concentrations were estimated using the Bradford assay.

### In vitro RNA synthesis

*In vitro* synthesized RNAs were made from double stranded DNA templates containing the T7ɸ2.5 +1A transcription start site RNA polymerase promoter (22). The templates were prepared by mixing equimolar amounts of single stranded template and non-template strands of DNA to a final concentration of 4 µM in 10 mM Tris-HCl at pH 8.0. The single stranded *in vitro* probe RNA was generated from DNA templates made from KD01 and KD02 (**Supplementary Table 3**). The double stranded RNAs (**Fig. 3B,C**) were made initially by generating single stranded RNAs of 25 and 40 nucleotides (nt), complementary to the probe RNA from (KD03, KD04) and (KD05, KD06) DNA templates, respectively (**Supplementary Table 3**). Next, the probe RNA was annealed with the 25 nt long RNA to generate 15 nt 5′ overhang RNA and 40 nt long RNA to generate blunt end RNA. Annealing of single stranded DNA or RNAs were performed in a thermocycler (Applied Biosystems, Pro flex PCR systems) by incubating the mixture at 80°C for 2 mins, cooling it to 37°C with a ramp rate set at 0.1°C per second, incubating for 10 mins, and finally incubating at 4°C for 10 min.

RNA products were generated using HiscribeT7 High Yield RNA Synthesis kit (New England Biolabs) as per manufacturer’s instructions. Briefly, 1 µM DNA template, 1 µl T7 RNA polymerase mix, 1X T7 Reaction Buffer were incubated with a final concentration of 7.5 mM of NTPs (ATP, GTP, UTP, CTP) in a 20 µL reaction volume at 37°C for 3 hours. For generating NAD^+^ capped RNAs, ATP was replaced with NAD^+^ (Sigma Aldrich) at a final concentration of 7.5 mM. Post incubation, DNase treatment was performed for 30 min at 37°C using 2U of TURBO DNase (Thermo Fisher Scientific) to degrade the template DNA. RNA was extracted post DNase treatment using acid phenol: chloroform: isoamyl alcohol, 25:24:1 (Sigma Aldrich) and recovered by ethanol precipitation. The RNAs were resuspended in RNase free water post recovery, and purity confirmed by TBE Urea PAGE gel electrophoresis on a 10% TBE Urea gel (Thermo fisher Scientific) and visualized by SYBR Gold staining (Thermo Fisher Scientific).

### 3′ Biotinylation of In vitro synthesized RNAs

*In vitro* synthesized single stranded probe RNA was subjected to 3′ biotinylation using pCp-Biotin (Jena Bioscience) in presence of T4 RNA ligase 1 (New England Biolabs). In a 30 µL reaction volume, RNAs up to 1 µg of RNA were incubated at 16°C overnight with 1,000 U T4 RNA ligase 1, in 1x T4 RNA ligase buffer, 10 mM ATP, 10% DMSO, 7.5% PEG8000, and 35 µM pCp-Biotin. The reaction mixture was incubated at 16°C overnight. Post incubation, the biotinylated RNAs were extracted using acid phenol: chloroform: isoamyl alcohol, 25:24:1 (Sigma Aldrich) and recovered by ethanol precipitation. Successful biotinylation at 3′ end of RNAs was verified by 10% TBE Urea PAGE gel electrophoresis followed by SYBR Gold staining (Thermo Fisher Scientific). Successful 3′ biotinylated RNAs migrate more slowly than non-biotinylated RNAs, indicated by a band shift.

### *E. coli* NAD cap RNA affinity purification (NcRAP)

NcRAP was performed as described previously for *S. cerevisiae* lysate (14). Briefly, three independent colonies (n=3; biological replicates) of *E. coli* K-12 strain MG1655 cultures grown overnight and each culture was diluted 100-fold in LB media and grown to an OD_600_ ≈ 1.2 at 37°C with shaking. The cells were then harvested by centrifugation and suspended in *E. coli* lysis reagent (New England Biolabs) in presence of 1 µL of T4 lysozyme (New England Biolabs) and 4U of TURBO DNase (Thermo Fisher Scientific) and incubated at room temperature for 10 min on a rocker. Post incubation, the cells were disrupted using Bioruptor (Diagenode; 5 cycles at high mode, 15 sec on 60 sec off) followed by centrifugation for 20 min at 16,000 x g. The supernatant containing bacterial cellular proteins was collected and mixed with 500 pmol of *in vitro* synthesized 5′ NAD^+^ capped or triphosphate end (5′ ppp) probe RNA immobilized on Dynabeads C1 Streptavidin Magnetic Beads (Thermo Fisher Scientific). RNA immobilization was done per manufacturer’s protocol and then incubated for 60 min on a rocker at 4°C. Free beads without immobilized RNA that were treated and washed the same to RNA bound beads served as a control to detect unspecific protein binding. The beads were immobilized on magnetic stands, washed 3 times with *E. coli* lysis reagent (New England Biolabs) supplemented with 60 mM NaCl to reduce nonspecific binding. The RNA bound proteins were eluted with the addition of 1x NuPAGE LDS sample buffer (Thermo Fisher Scientific), followed by heating the bead bound RNA-protein complexes at 95°C for 5 min. Eluates were resolved by electrophoresis on a 4-12% NuPAGE Bis-Tris mini protein gel (Thermo Fisher Scientific) and subsequently visualized (**Fig. 1A**) by staining with SimplyBlue safe stain (Thermo Fisher Scientific). The entire eluate from NAD^+^ capped RNA, 5′ ppp RNA, and the beads alone were sent for mass spectrometry analysis (**Supplementary Table 4**). Additionally, we excised the prominent band at ∼70 kDa (**Fig. 1A**) and analysed it as described below for mass spectrometry analysis (**Supplementary Table 5**).

**Figure 1.**
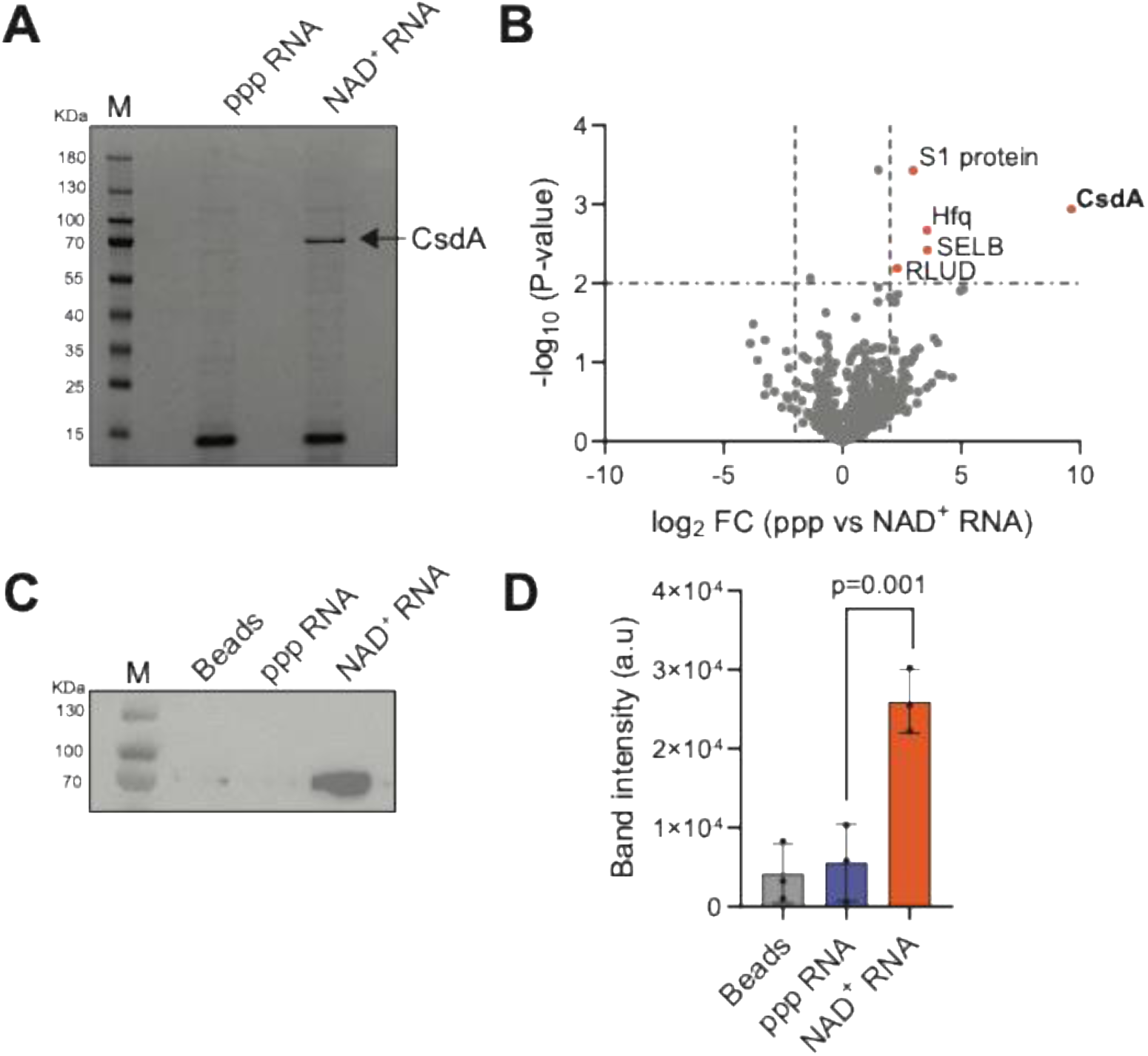
Identification of CsdA as an NAD^+^ RNA interacting protein by NAD cap RNA affinity purification using *E. coli* MG1655 lysate (**A**) SDS PAGE analysis of proteins interacting with NAD^+^ capped or ppp RNA after elution in NcRAP (**B**) Volcano plot of proteins differentially enriched in NAD^+^ capped vs triphosphate (ppp) RNA fractions. The x-axis corresponds to log_2_ fold change of protein association (NAD^+^ cap vs ppp RNA); while the y-axis shows -log_10_ transformed P values. Dashed lines indicate significance threshold defined by log_2_ fold change > 2 and -log_10_ P value > 2; proteins meeting both criteria were considered significantly enriched. A two-sided t-test was applied for comparison. (**C**) Western blot analysis of NcRAP performed with *E. coli* MG1655 c*sdA-3xFLAG* strain. CsdA levels were detected using anti-FLAG antibody. (**D**) Quantification of western blot shown in (C). Data represent three independent biological replicates. Points indicate mean values, and error bars represent standard deviation. Statistical analysis was performed using a two-tailed t-test.

### Proteomics sample preparation and liquid Chromatography-tandem mass spectrometry (LC-MS/MS)

Protein complexes after affinity purification were processed using E3filters, following the procedure as described previously (23). Briefly, the protein eluants were mixed with an equal volume of SDS buffer (5% SDS, 100 mM Tris-HCl, pH 8) and a final concentration of 10 mM tris(2-carboxyethyl)phosphine (TCEP) and 40 mM 2-chloroacetamide (CAA), and boiled at 95°C for 10 min. Afterward, the samples were mixed with a 4x volume of 80% acetonitrile (ACN) and then transferred to E3filters (CDS Analytical, Oxford, PA). The filters were spun at 3,000 rpm for 2 min and washed twice with 400 μL of 80% ACN. For digestion, 200 μL of 50 mM triethylammonium bicarbonate (TEAB) buffer with 1 μg of trypsin/Lys-C mix was added to the samples and then incubated at 37°C for 16-18 h with gentle shaking. After digestion, peptides were eluted sequentially with 200 μL of 0.2% formic acid (FA) in water, and 200 μL of 0.2% FA in 50% CAN. The peptides were dried in SpeedVac and desalted using C18 based StageTips (CDS Analytical).

The LC-MS/MS analysis was performed following the protocol as described previously with minor changes (23). The LC separation was performed on an Ultimate 3000 RSLCnano system coupled with a trap column (PepMap100 C18, 300 μm × 2 mm, 5 μm) and an analytical column (PepMap100 C18, 15 cm × 75 μm i.d., 3 μm). Mobile phase A was 0.1% formic acid (FA) in water; mobile phase B was 0.1% FA in acetonitrile. A linear LC gradient was applied from 1% to 25% mobile phase B over 125 min, followed by an increase to 32% mobile phase B over 10 min. The column was washed with 80% mobile phase B for 5 min, followed by equilibration with mobile phase A for 15 min. The MS analysis was performed on an Orbitrap Eclipse system installed with field asymmetric ion mobility spectrometry (FAIMS) Pro Interface (Thermo Fisher Scientific). For the ion source settings, the spray voltage was set to 1.7 kV, funnel RF level at 30%, and heated capillary temperature at 275°C. The MS data were acquired in Orbitrap at 60,000 resolution, followed by data dependent acquisition (DDA) MS/MS of the most intense precursors for 1 second. The MS1 scan range was set to 375–1,500 m/z, with automatic gain control (AGC) target set to 4E5, and the maximum injection time mode was set to Auto. For MS2 analysis, precursors with charge states 2–5 were selected. The isolation mode was Quadrupole, collision was by HCD at 30% normalized collision energy (NCE). The Orbitrap was set to detect MS2 fragments at 15,000 resolution; AGC target was set to 5E4, and maximum injection time was 22 ms. Monoisotopic precursor selection (MIPS) was set to Peptide. For FAIMS settings, a 3-CV experiment (−40|-55|-75) was applied.

The (DDA) MS data were processed using MaxQuant and Andromeda software suite (version 2.5.2.0) and were searched against the *E. coli* database obtained from UniProt Knowledgebase (K12 strain; 4,598 sequences). The enzyme specificity was set to ’Trypsin’. The variable modifications included oxidation of methionine, and acetyl (protein N-terminus). The fixed modification included carbamidomethylation of cysteine. The minimum number of amino acids required for peptide identification was 7. The false discovery rate (FDR) was set to 1% for protein and peptide identifications. MaxLFQ function embedded in MaxQuant was enabled for label-free quantitation, and the LFQ minimum ratio count was set to 1. Differential protein abundance was obtained using a two-sided t-test in Perseus software (version 1.6.2.3) (24), and proteins with log_2_ fold change > 2 and -log_10_ p-value > 2 were considered to be significant.

### RNA Extraction

Total cell RNA was extracted from four independent *E. coli* MG1655 cultures (n=4) grown to OD_600_ ≈ 1.2. The cultures were flash frozen in dry ice and stored in - 80°C before RNA extraction. Initially, the frozen cell pellet was resuspended in 1ml TRI reagent (Molecular Research Center), heated at 65°C for 10 min, and centrifuged for 10 min at 15,000 g at room temperature to pellet insoluble materials. The supernatant was collected, and total RNA was purified using the Direct-zol RNA MiniPrep Plus kit (Zymo Research) according to manufacturer’s instructions. Post elution, a DNase treatment step was performed by incubating 10 µg of RNA with 2U of Turbo DNase (Thermo Fisher Scientific) for 30 min at 37°C. Post incubation, RNAs were extracted using acid phenol: chloroform: isoamyl alcohol, 25:24:1 (Sigma Aldrich) and recovered by ethanol precipitation.

### NudC/NudC E178Q treatment

Total cellular RNAs extracted from *E. coli* MG1655 cells were treated with either 1 µM of NudC, NudC E178Q, or mock treated in the presence of NEBuffer 2.1 (New England Biolabs) at 37°C for 1 hour and then purified using Direct-zol RNA MiniPrep Plus kit (Zymo Research) according to manufacturer’s instructions (**Fig. 3D**).

### Electrophoretic Mobility Shift assay (EMSA)

Purified CsdA of varying concentrations was incubated with 100 nM of *in vitro* synthesized 5′ NAD^+^ capped or ppp probe RNA at 37°C for 30 min in EMSA buffer containing 25 mM Tris, 100 mM KCl, 5 mM MgCl_2_, 10% glycerol, 1 mM DTT, and 0.1 mM EDTA (**Fig. 2A**). RNA-protein complexes were resolved by electrophoresis on a 5% native polyacrylamide gel, run in 0.5x TBE at 180 V for 35 min at room temperature. The gel was stained with SYBR Gold nucleic acid gel stain (Thermo Fisher Scientific) and visualized using ChemiDoc imaging system (Bio-Rad). Image quantification was done using Image Lab software (Bio-Rad). This assay was performed in three (n=3) independent technical replicates. Statistical analysis was performed by comparing the means using one-sided Welch’s t-test only at the highest concentration of CsdA (0.7 µM).

**Figure 2.**
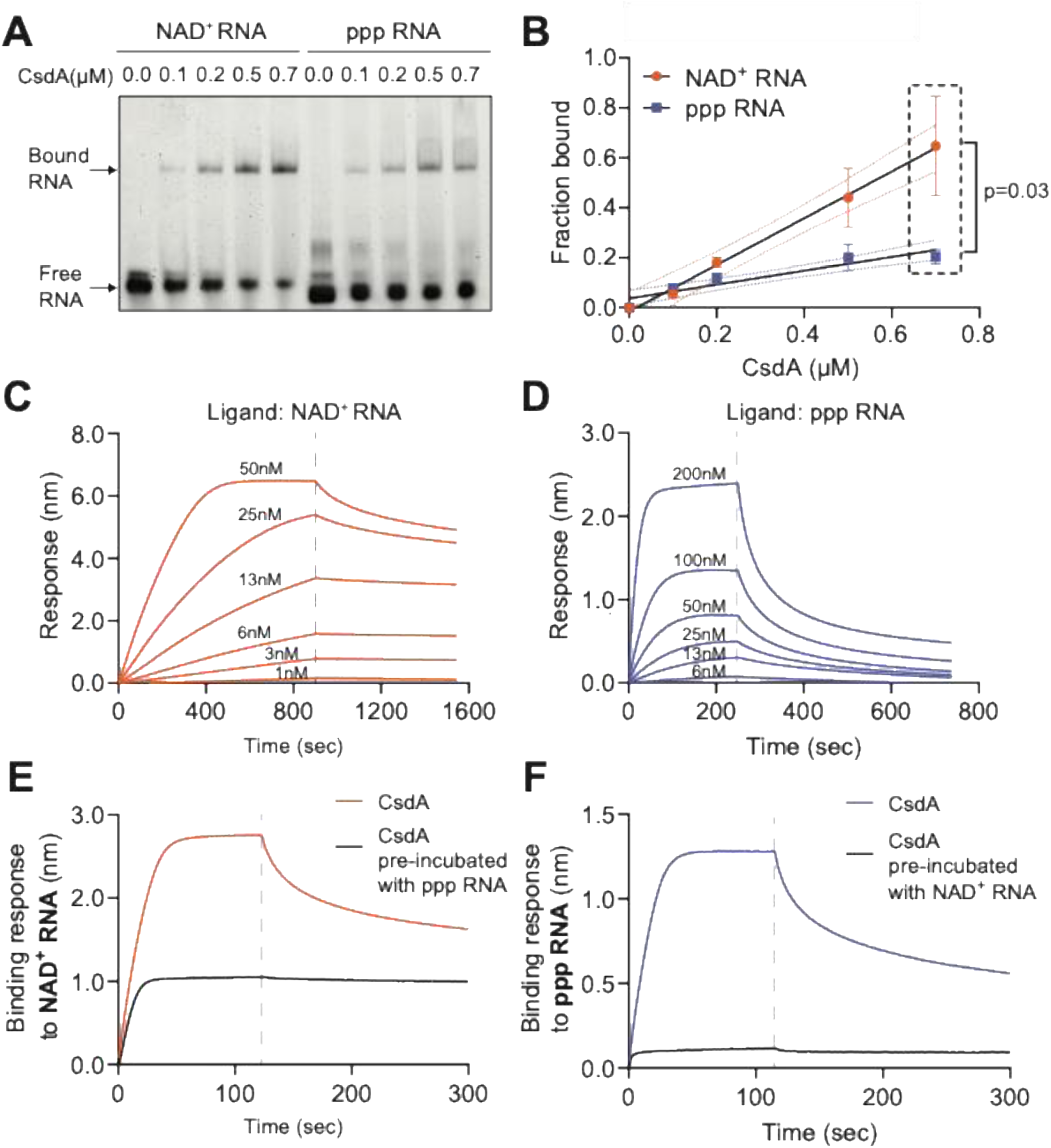
Biochemical characterization of CsdA interaction with NAD^+^-capped RNA. (**A**) Electrophoretic mobility shift assay (EMSA) of CsdA binding to 5′ NAD^+^ capped or 5′ ppp probe RNA. Gel stained with SYBR Gold nucleic acid stain. (**B**) Quantification of CsdA–RNA binding in (A). Data represent three independent replicates. Points indicate mean values, and error bars represent standard deviation. Welch’s one-sided *t*-test was performed for the highest CsdA concentration (0.7 µM). (**C,D**) Biolayer interferometry (BLI) analysis using purified CsdA and immobilized (**C**) NAD^+^-capped or (**D**) ppp probe RNA. Sensograms of biosensor responses over time for increasing concentrations of CsdA interacting with a fixed concentration (50 nM) of immobilized RNA. Experiments were performed in triplicate. (**E**) RNA binding competition assays. Biosensor responses for 100 nM CsdA alone or 100 nM CsdA pre-incubated with 200 nM ppp RNA for 10 min at 37°C prior to interaction with 50 nM immobilized NAD^+^-capped RNA. (**F**) Biosensor responses for 100 nM CsdA alone or 100 nM CsdA pre-incubated with 200 nM NAD^+^-capped RNA for 10 min at 37°C prior to interaction with 50 nM immobilized ppp RNA.

### Biolayer interferometry

Biolayer interferometry was performed to provide a more detailed insight into CsdA interaction with 5′ NAD^+^ capped vs triphosphate (ppp) RNA. The assay was carried out on an Octet RH16 instrument (Sartorius) (25). 3′ biotinylated *in vitro* synthesized 5′ NAD^+^ or ppp probe RNA was immobilized on Octet streptavidin (SA) biosensors (Sartorius), equilibrated with binding buffer (50 mM Tris, 200 mM NaCl, 0.04% Tween 20, and 1.5 mM ATP), followed by blocking the free streptavidin sites with a final concentration of 10 µg/mL biocytin (Sigma Aldrich) before introduction into binding buffer containing varying concentrations of CsdA (as indicated in results). Raw data were extracted, analysed using Octet Data Analysis HT Software (version 9.0), and responses (nm) were plotted in GraphPad Prism (**Fig. 2C, D**). The kinetic parameters (**Table 1**) were estimated from three independent (n=3) technical replicates.

**Table 1.** Binding parameters for the interaction between CsdA and RNA.

| | $K_D$ (nM) | $k_{on}$ (M<sup>-1</sup>s<sup>-1</sup>) | $k_{off}$ (s<sup>-1</sup>) |
| --- | --- | --- | --- |
| 5' NAD<sup>+</sup> RNA | 1.6±0.7 | (7.9±2)*10<sup>4</sup> | (1.5±1)*10<sup>-4</sup> |
| 5' ppp RNA | 13±2.1 | (2.3±0.7)*10<sup>5</sup> | (2.8±0.8)*10<sup>-3</sup> |

For the competition assay, 100 nM CsdA was incubated with 200 nM non-biotinylated *in vitro* synthesized 5′ ppp probe RNA for 10 min at 37°C in one of the 96 wells before the introduction of streptavidin biosensor having immobilized NAD^+^ capped probe RNA (**Fig. 2E**). Conversely, CsdA was incubated with non-biotinylated *in vitro* synthesized NAD^+^ capped probe RNA for 10 min at 37°C before the introduction of immobilized 5′ ppp probe RNA (**Fig. 2F**). Raw data was processed using Octet Data Analysis HT Software (version 9.0), extracted, and responses (nm) were plotted in GraphPad Prism.

### ATPase assay

ATPase activity of CsdA was measured with *in vitro* synthesized single stranded or double stranded RNA having a 5′ NAD cap or ppp in presence of 100 nM of purified protein (**Fig. 3A-C**). To assess the ATPase activity of CsdA in presence of whole cell RNAs, 200 nM of CsdA was incubated with 500 ng of total RNA from untreated, NudC treated, and NudC E178Q treated samples (**Fig. 3D**). The reaction was carried out at 25°C, supplemented with NADH regeneration system (pyruvate kinase/lactate dehydrogenase (Sigma Aldrich), 10 mM NADH, 50 mM phosphoenolpyruvate, and 10 mM ATP). The study was performed in a clear 384-well plate with clear flat bottoms (Corning). Briefly, hydrolysis of ATP by CsdA generates ADP, which is utilized by pyruvate kinase (PK) to form pyruvate, which is reduced to lactate by lactate dehydrogenase (LDH), simultaneously oxidizing NADH to NAD^+^. The oxidation of NADH was measured over time by following loss of 340 nm absorbance. The reaction buffer consisted of 25 mM Tris, 100 mM KCl, 5 mM MgCl_2_, 1 mM DTT, 0.1 mM EDTA, and 10% glycerol. Reactions were monitored using Magellan Data analysis software (TECAN Life Sciences). ATPase rate was quantified by determining the linear rate of NADH oxidation over time. For the *in vitro* synthesized RNA experiments (**Fig. 3A-C**), each RNA concentration was measured in triplicate (n=3) for all 3 substrate types. For the total cell RNA experiment (**Fig. 3D**), four independent biological replicates (n=4) for each treatment condition were used. Statistical analysis was performed by comparing the means using a one-sided t-test.

**Figure 3.**
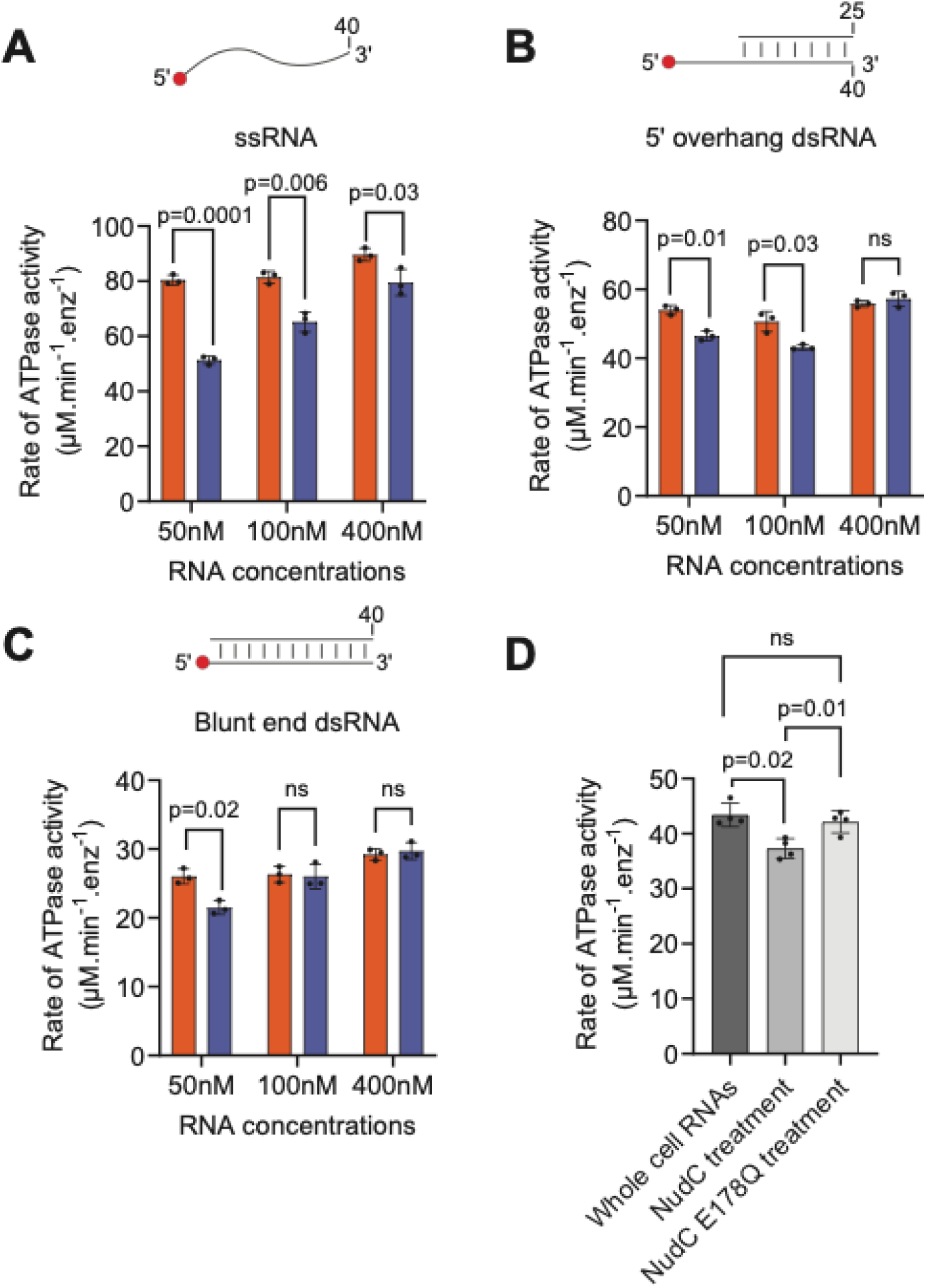
5′ NAD^+^ capping influences CsdA ATPase activity. (**A**, **B**, and **C**) Rates of CsdA ATPase activity in presence of RNA substrates bearing either a 5′ NAD^+^ cap (orange) or ppp end (blue). ATPase rate of 100 nM CsdA in presence of varying amounts of RNA as indicated over time at 25°C. with ssRNAs (**A**), double stranded 5′-overhang RNA (**B**), and double-stranded blunt end RNA (**C**). Data represent three independent replicates. Points indicate mean values, and error bars represent standard deviation. Statistical significance was assessed using multiple unpaired t-tests; adjusted P values are reported above the bars. (**D**) ATPase activity of 200 nM CsdA in the presence of 500 ng of *E. coli* whole cell RNA, which were mock treated, NudC treated, or inactive mutant NudC E178Q treated. Data represent four independent replicates. Points indicate mean values, and error bars represent standard deviation. Statistical analysis was determined using a one-sided t-test.

### Western Blot

*E. coli* MG1655 csdA-*3xFLAG* strain was grown overnight in LB media at 37°C with shaking. Next day, cultures were back diluted 1:100 in fresh media and grown to OD_600_ ≈ 1.2. Cell lysates were generated and incubated with streptavidin bead-bound RNAs as mentioned in *E. coli* NcRAP method above. To elute protein from the pulldowns, the beads were boiled with SED buffer (4% SDS, 10 mM EDTA, 100 mM Tris HCl, 20 mM DTT, 0.05% Tween 20, pH 8.0). All pulldown samples were quantified using A_280_ readings on a DS-11 spectrophotometer (Denovix). Equal amount of protein (30 µg) was loaded onto a NuPAGE Bis-Tris Mini protein gel 4-12% (Thermo Fisher Scientific) for all samples and run with 1X MOPS buffer. Protein samples were transferred onto a nitrocellulose membrane (Bio-Rad) using a semidry transfer blot apparatus (Bio-Rad). To detect CsdA-3xFLAG protein, the membrane was probed with primary M2 anti-FLAG mouse monoclonal antibodies (1:500 dilution) (Sigma Aldrich) and anti-mouse HRP secondary antibodies (1:10,000 dilution) (Thermo Fisher Scientific). The membrane was developed with Clarity western ECL substrate (Bio-Rad) and visualized using ChemiDoc imaging system (Bio-Rad). Band intensity was quantified using ImageJ software (version 1.54r) and plotted using GraphPad Prism. Three independent biological replicates were used for this study. Statistical analysis was performed by comparing the means using a two-sided t-test.

### DIC imaging of RNA-dependent CsdA phase separation

DIC imaging samples (10 µL) were prepared by combining components in the following order: 20 mM TrisCl pH 7.0, 100 mM NaCl, 15 mM ATP, 5 µM CsdA, 250 µM MgCl_2_, and 200 nM *in vitro* synthesized NAD^+^ or ppp probe RNA (**Fig. 4A, F, G**). To test salt response or RNase A treatment, the samples were prepared in the same order with 1 M NaCl added instead of 100 mM NaCl. At room temperature, RNase A (1 µg) was added last using the same order and conditions stated previously, with the samples inverted for thirty minutes before imaging. For imaging, 8 µL of sample was pipetted onto the glass slide with an adhesive well, the coverslip was placed and then inverted for approximately 30 min to allow for CsdA condensate wetting. Images were taken on a Nikon TiE inverted setup using an APO Lambda 100X oil objective with differential interference contrast (DIC) using Zeiss Immersol 518F immersion oil. Additionally, the Nikon Element AR software was used to control the imaging setup, and images were collected using an Andor Ixon Ultra 897 EMCCD camera. DIC imaging uses a total polarizer until an equal visual ratio of the dark and light sides of the image is present. Statistical analyses for size of condensates and number of condensates per µm2 was performed by comparing the means using one-sided Welch’s t-test (**Fig. 4B, C**).

**Figure 4.**
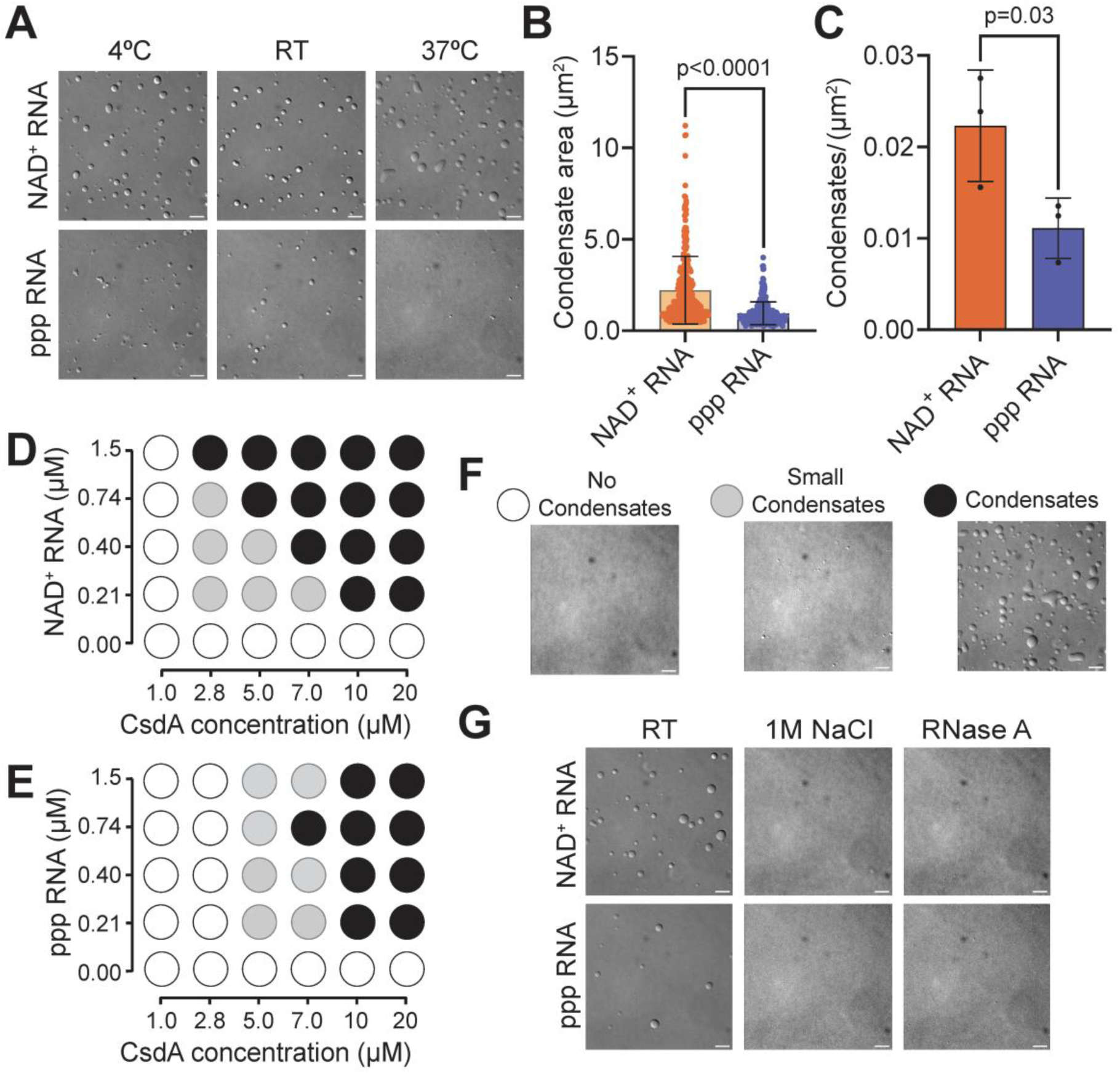
5′ NAD caps influence biomolecular condensate formation by CsdA. (**A**) *In vitro* phase separation of 5 µM CsdA and 200 nM NAD^+^ capped or ppp probe RNA at 16°C, room temperature (RT) or 37°C as visualized by DIC microscopy. Images representative of at least three independent experiments. Scale bar = 5µm. (**B**) Quantitative analysis of condensate area at RT. Mean values calculated from three independent replicates (n ≥ 300). Bars indicate mean values, and error bars denote standard deviation. Statistical significance determined using Welch’s one-sided t-test. (**C**) Quantification of the average number of condensates formed per µm^2^. Each Dot indicates the total number of droplets per image per pixel area (µm^2^). Total number of droplets counted for each replicate ≥ 300. Bars indicate mean values, and error bars denote standard deviation. Statistical significance assessed using Welch’s one-sided t-test. (**D, E**) Phase diagrams of purified CsdA mixed with (**D**) NAD^+^ capped or (**E**) ppp RNA in varying concentrations. (**F**) Three distinct regimes are observed in condensate formation assays as visualized by white (No condensates), grey (Small condensates) or black spheres (Large condensates). (**G**) Formation of phase-separated condensates by CsdA are dependent on ionic interaction between macromolecules and RNA presence as shown by treatment with 1M NaCl or 1µg RNase A.

### *In vitro* NAD^+^ RNA decapping assay using NudC

*In vitro* NAD^+^ RNA decapping assay using NudC was performed as described previously (26). Substrate NAD^+^ capped RNAs were generated by *in vitro* transcription. 100 units of T7 RNA polymerase (New England Biolabs) and 0.6 µM DNA template (same as used to generate probe RNA) were incubated in 1x RNA Pol Reaction Buffer (New England Biolabs) in presence of 1 mM NAD^+^ along with 200 µM of GTP, CTP, and UTP, supplemented with 5 µCi [⍺32-P] radiolabeled GTP and incubated for 30 min at 37°C. The reaction was terminated using stop solution (20 mM Tris, pH 8.0, 50 mM sodium acetate, and 0.1 mg/mL glycogen), followed by RNA extraction using acid phenol: chloroform: isoamyl alcohol, 24:25:1 (Sigma Aldrich) and ethanol precipitation. RNA pellets were suspended in NEBuffer 2.1 (New England Biolabs) and were incubated with 1 µM NudC or NudC storage buffer, with or without an indicated amount of CsdA. Reactions were incubated for 15 min at 37°C and were terminated by the addition of RNA loading dye (100 mM Tris HCl, pH 8.0, 90% Formamide, 18 mM EDTA, 0.025% bromophenol blue, 0.025% xylene cyanol). The RNAs were resolved by PAGE gel electrophoresis by running the samples on a 10% TBE urea polyacrylamide sequencing gel (National Diagnostics). Gels were exposed to phosphorimaging screens (Cytiva) and imaged using a Sapphire Biomolecular Imager (Azure Systems). Bands were quantified using Image Lab software (Bio-Rad) and plotted using GraphPad Prism (Dotmatics). Statistical analysis was performed using simple linear regression to get the best-fit line.

### Statistical Analysis

For Figure 1, statistical significance for both datasets (**Fig. 1B, D**) was assessed using a two-sided t-test, as no a priori directional hypothesis was defined. However, subsequent comparisons (**Fig. 2 through 4**) of means between datasets were performed using a one-sided t-test, based on the directional hypothesis that 5′ NAD^+^ capping on RNA enhances CsdA binding relative to 5′ ppp RNA, consistent with trends observed in Figure 1. All statistical analysis were performed using GraphPad Prism software (version 10.6; Dotmatics).

## RESULTS

### Identification of *E. coli* CsdA as an NAD^+^ cap RNA binding protein

As a first step in elucidating the functional consequences of NAD^+^ capping, we set out to identify *E. coli* proteins that bind specifically to NAD^+^ capped transcripts using NAD cap RNA affinity purification (NcRAP), a method initially used to identify novel NAD^+^ decapping enzymes in *S. cerevisiae* (14). A single stranded *in vitro* probe RNA incorporating a 5′ NAD^+^ cap (NAD^+^ RNA) and 3′ biotin served as bait, while a 5′ triphosphate version of the same RNA (ppp RNA) was used as the biological control. The probe RNA sequence was specifically selected as it generated no secondary structure (14). We performed NcRAP pulldowns with NAD^+^ RNA, ppp RNA, or streptavidin beads alone using *E. coli* MG1655 cell lysate and analysed the full set of captured proteins by LC-MS/MS (**Fig. 1**). Captured proteins were resolved by SDS-PAGE (**Fig. 1A**). Notably, a prominent ∼70 kDa band was clearly present in the NAD^+^ RNA pulldown, but absent from the ppp RNA pulldown. Analysis of the excised band by liquid chromatography-tandem mass spectrometry (LC-MS/MS) -based quantitative proteomics identified the band as the *E. coli* RNA helicase CsdA (**Supplementary Table 5**). Interestingly, ribosomal protein S1 was detected as a prominent NAD^+^ RNA interactor second only to CsdA in the MS analysis (**Supplementary Table 4**). Comparison of proteins differentially captured by NAD^+^ RNA and ppp RNA (or streptavidin beads alone) revealed that CsdA was enriched in the NAD^+^ RNA bound samples, indicating its preference for NAD^+^ capped RNAs (**Fig. 1B**, **Supplementary Table 4**). Five proteins were enriched at least 5-fold in the NAD^+^ RNA pulldown compared to the ppp RNA dataset, with p-values below 0.01 (**Fig. 1B**). Of these, the most highly enriched was CsdA, which was ∼1000-fold more abundant in the NAD^+^ RNA dataset, corroborating our SDS-PAGE analysis.

In addition to CsdA, two well-studied single stranded RNA interacting (27,28) proteins were strongly enriched in the NAD^+^ RNA over ppp RNA datasets: the small regulatory RNA chaperone Hfq (12-fold) and ribosomal protein S1 (8-fold). Hfq is one of the primary chaperones of small regulatory RNAs (sRNAs) in *E. coli*, many of which have been reported to carry NAD^+^ caps (^1,4^). Ribosomal protein S1 is an essential part of the 30S ribosomal subunit, which interacts with the 5′ untranslated region of RNA adjacent to the Shine Dalgarno element and acts as an RNA chaperone, unfolding highly structured mRNA leaders (29). Our results implicate CsdA, Hfq and ribosomal protein S1 as having strong binding preferences for 5′ NAD^+^ capped RNAs *in vivo*.

Interestingly, we failed to detect any significantly enriched RNA binding proteins in the ppp RNA dataset relative to NAD^+^ RNA dataset. The lack of strong ppp RNA-specific binders may be due to the specific sequence of the *in vitro* transcribed probe RNA used in our pulldown, the absence of 5′ end secondary structures (30), or may indicate that binding partners bind less tightly to triphosphate 5′ ends than to NAD^+^ caps.

Given the exceptional enrichment of CsdA in the NAD^+^ RNA pulldown, we chose to further characterize the interaction between CsdA and NAD^+^ RNA. We repeated the NcRAP method using an *E. coli* strain containing a modified *csdA* gene with a C-terminal 3xFLAG epitope tag. Western blot analysis of pulldown eluates post NcRAP showed significant enrichment of CsdA-3xFLAG in the NAD^+^ RNA pulldown compared to pulldown with ppp RNA or the streptavidin beads alone (**Fig. 1C, D**). Taken together, these findings validate CsdA as a prominent and highly selective binder of 5′ NAD^+^ capped RNAs in *E. coli*.

### Biochemical characterization of NAD^+^ cap interaction with CsdA

We next sought to assess the binding preference of CsdA for NAD^+^ capped RNAs over ppp RNAs *in vitro*. First, we performed electrophoretic mobility shift assays (EMSAs) that followed the migration of *in vitro* NAD^+^ or ppp RNA probes binding in the presence of increasing concentrations of CsdA (**Fig. 2A**). CsdA formed a stable RNA protein complex with both RNAs, evidenced by the appearance of a single band with slower electrophoretic mobility. However, significantly more RNA protein complex formed with NAD^+^ RNA than with ppp RNA, especially at the highest CsdA concentration (**Fig. 2B**). This indicates preferential binding of CsdA to NAD^+^ RNA, consistent with our NcRAP results.

To quantitatively assess RNA binding by CsdA, we monitored the interaction of purified CsdA with immobilized NAD^+^ RNA or ppp probe RNA of the same sequence used in NcRAP by biolayer interferometry (BLI) over a range of protein concentrations (**Fig. 2C, D**). We found that CdsA binds with an 8-fold lower equilibrium dissociation constant (*K*_D_) to NAD^+^ RNA (1.6 nM) than to ppp RNA (13 nM) (**Table 1**). CsdA also exhibited slower binding kinetics to NAD^+^ RNA than ppp RNA, with a ∼3-fold slower association rate constant (*k*_on_) and a ∼19-fold slower dissociation rate constant (*k*_off_). The slower *k*_off_ value suggests that CsdA makes more stable contacts with NAD^+^ RNA. We also noted that saturating concentrations of CsdA resulted in ∼2.5-fold higher equilibrium binding response to immobilized NAD^+^ RNA than to ppp RNA, suggesting higher binding stoichiometry to NAD^+^ RNA. In addition, we did not observe any significant influence of 50,000 molar excess of free NAD^+^ on the binding efficiency of CsdA with NAD^+^ RNA (**Supplementary Fig. 2A**), indicating NAD^+^ RNA as the preferred substrate versus free NAD^+^.

Next, we tested a more physiologically relevant scenario by assessing how the presence of an RNA competitor in solution influences CsdA binding to immobilized RNA. We pre-incubated 100 nM of CsdA with 200 nM of RNA competitor for 10 min, then monitored binding to immobilized RNA by BLI. Whereas CsdA binding to immobilized NAD^+^ RNA resulting in a binding response of ∼2.6 nm in the absence of competitor, the presence of ppp RNA competitor reduced the binding response by ∼60% to ∼1 nm, indicating moderate competition (**Fig. 2E**). By contrast, strong competition was observed in the inverse experiment: CsdA binding to immobilized ppp RNA resulted in ∼1.1 nm binding response in the absence of competitor, while the presence of NAD^+^ RNA competitor reduced this value by ∼90% to ∼0.1 nm (**Fig. 2F**). This result demonstrates that CsdA retains a strong preference for NAD^+^ capped RNAs when presented with a mixture of RNA species. Interestingly, we observed that CsdA binding to ppp RNA was more sensitive to salt concentrations than binding to NAD^+^ capped RNA (**Supplementary Fig. 2B**), which indicates that binding between CsdA and 5′ NAD cap is mediated to a greater degree by non-electrostatic forces, such as hydrophobic or base stacking interactions. Collectively, the EMSA and BLI data demonstrate that CsdA exhibits strong binding specificity and suggest a distinct mode of interaction with NAD^+^ capped RNA over ppp RNA.

### NAD^+^ capped RNA influences CsdA ATPase activity

We next assessed whether the preferential binding of CsdA to NAD^+^ RNA affects one of its native functions. CsdA possesses a low basal rate of ATP hydrolysis, which is strongly stimulated upon interaction with RNA (31).The ATPase activity of CsdA is necessary for its helicase function and is required for optimal growth of *E. coli* at low temperatures (31,32).

We evaluated the effects of NAD^+^ RNAs and ppp RNAs on the ATPase activity of purified CsdA. Three structurally distinct RNAs were generated: the 40 nucleotide single stranded RNA without predicted secondary structure used in previous experiments (ssRNA), a double stranded RNA containing the 40 nucleotide RNA bound to a complimentary strand with a 15 nt overhang (5′ overhang dsRNA), and a 40 nucleotide double stranded RNA with blunt ends (Blunt end dsRNA). Versions of each RNA were prepared bearing either a 5′ NAD^+^ cap or a 5′ ppp end. The ATPase activity of CsdA was measured in the presence of each RNA construct. Considering the low physiological abundance of NAD^+^ capped RNAs in *E. coli* whole cell RNA (1) we kept a fixed concentration of CsdA and varied RNA concentrations from 50 nM to 400 nM. CsdA exhibited the highest ATPase activity in the presence of ssRNA constructs (**Fig. 3A**), with 5′ overhang dsRNA stimulating intermediate activity (two-thirds of the ssRNA level; **Fig. 3B**), and blunt end dsRNA stimulating the lowest activity levels (one-third of the ssRNA level; **Fig. 3C**). This behaviour is in line with prior observations (31) and suggests that an accessible single-stranded overhang aids CsdA engagement. Importantly, we observed that NAD^+^ RNAs stimulated higher ATPase activity than ppp RNAs, although the magnitude of this difference depended on the specific RNA construct and concentration.

We observed the strongest rate of ATP hydrolysis by CsdA in presence of ssRNA (**Fig. 3A**). At 50 nM ssRNA, we measured 1.6-fold higher ATPase activity in the presence of NAD^+^ ssRNA than ppp ssRNA. This trend persisted at higher ssRNA concentrations, although the differential decreased to 1.1-fold at 400 nM ssRNA. NAD^+^ capped dsRNAs also stimulated higher ATPase levels than ppp dsRNAs at low concentrations (∼1.2-fold), although at the highest dsRNA concentrations we observed no significant difference in ATPase rate between differentially capped dsRNA. Thus, while high levels of both NAD^+^-capped and ppp RNAs can produce similar stimulatory effects on CsdA’s ATPase activity, its tighter binding to NAD^+^ capped RNAs produces equivalent stimulation even at sub-saturating levels of these RNAs. Furthermore, we did not observe any significant influence of free NAD^+^ on ATPase rate of CsdA (**Supplementary Fig. 3A**).

To better elucidate the biochemical activity of CsdA, we employed total cellular RNA to more closely mimic its native intracellular environment. Whole cell RNAs from *E. coli* MG1655 cells were treated with NudC to remove 5′ NAD^+^ caps, treated with the enzymatically inactive NudC E178Q variant, or mock treated. We evaluated the ATPase activity of CsdA in presence of each RNA sample and found that whole cell RNA stimulated ATPase activity of CsdA (**Fig. 3D**), as observed with *in vitro* synthesized RNA. Importantly, isolated total cellular RNA that was subsequently NudC-treated stimulated CsdA ATPase activity to a ∼14% lower level than mock-treated RNA. By contrast, treatment of RNA with NudC E178Q did not significantly alter the ATPase activity of CsdA compared to mock-treated RNAs. This behaviour is consistent with a stronger stimulatory effect by NAD^+^-capped RNA. Moreover, these findings suggest that lower-abundant NAD^+^ capped transcripts *in vivo* can effectively modulate CsdA ATPase activity and may consequently influence the helicase activity of CsdA.

### 5′ NAD+ cap RNA influences biomolecular condensate formation of CsdA

CsdA is a member of the RNA dependent DEAD box family of ATPases, which has been shown to undergo phase separation to form membrane-less compartments called RNP condensates (33). In a recent study, *in vivo* condensate formation by CsdA was shown to help *E. coli* shift from proliferation to dormant state under nutrient limiting conditions (34). Additionally, CsdA was reported to be indispensable for growth of *E. coli* at low temperatures (15,16). Given our observation of the positive influence of NAD^+^ cap on ATPase activity of CsdA (**Fig. 3**), we wanted to understand the influence of 5′ NAD^+^ capping and temperature on condensate formation by CsdA. To that end, we incubated purified CsdA with *in vitro* synthesized NAD^+^ or ppp probe RNA for 30 min and observed condensate formation under a range of temperatures. We found that NAD^+^ RNA stimulated CsdA phase separation across the full 4 to 37°C range tested. In contrast, ppp RNA formed smaller condensates only at 4°C and room temperature, with phase separation severely attenuated at 37°C (**Fig. 4A**). Importantly, quantitative analysis at room temperature (RT) conditions showed an increase in both size and number of condensates formed by CsdA in presence of NAD^+^ RNA compared to ppp RNA. Comparison of condensates at RT, we observed ppp RNA produced condensates with an average area of 1.3 µm² and a density of 0.011 condensates/µm², whereas NAD^+^ RNA yielded larger (2.3 µm²) and more abundant condensates (0.022 condensates/µm²) (**Fig. 4B, C**).

To better understand how RNA and CsdA concentration influence condensate formation, we tested an array of concentrations and categorized the results into three regimes: no assembly, small condensates (diameter < 0.5 µm), and large condensates (diameter 1.5– 7.5 µm) (**Fig. 4F**). The results were plotted as 2-dimensional phase diagrams. NAD^+^ RNA shifted the phase boundary toward lower CsdA and RNA concentrations, reducing the critical saturation concentration (*C*_sat_) to approximately 2.8 µM across all tested RNA concentrations. By contrast, the phase boundary was essentially constant across the range of tested ppp RNA concentrations, with a *C*_sat_ of 5 µM (**Fig. 4D, E**). Additionally, to exclude the possibility that free NAD^+^ independently influences condensate formation, CsdA was incubated with 5 mM NAD^+^, and condensate formation was assessed under similar conditions. At 5 µM and 10 µM CsdA concentration, only small condensates (diameter < 0.5 µm) were observed in low abundance, with no large condensate formation (**Supplementary Fig. 3B**). These results indicate that *in vitro* phase separation by CsdA into condensates is specifically enhanced by NAD^+^ capped RNA, but not by free NAD^+^. Furthermore, to determine whether CsdA condensate formation depends on electrostatic interactions and on RNA, we treated co-assemblies with 1 M NaCl or 1 µg RNase A. Addition of either 1 M NaCl or 1 µg RNase completely dissolved CsdA condensates regardless of RNA cap status, confirming that condensate formation requires both electrostatic and RNA-mediated interactions (**Fig. 4E**). These results demonstrate that 5′ NAD^+^ capping on transcripts influences RNA-dependent condensate formation by CsdA.

### CsdA accelerates deNADing activity of NudC

A recent study demonstrated by protein pulldown and bacterial two-hybrid analysis that CsdA and NudC physically associate (19). As both proteins are NAD^+^ capped RNA interactors, we set out to test if CsdA influences the NAD^+^ RNA decapping activity of NudC. For *in vitro* decapping assays, we chose a NudC concentration (0.5 μM) that resulted in cleavage of approximately 50% of NAD^+^ capped RNA substrate (**Supplementary Fig. 4A, B**). NudC dependent decapping of NAD^+^ RNA was assayed across a range of CsdA concentrations. We observed that CsdA dose-dependently stimulated NudC activity on NAD^+^ capped transcripts, resulting in an increase in decapping from ∼30% in the absence of CsdA to ∼80% in the presence of 2.6-fold excess of CsdA of NudC. (**Fig. 5A, B**). We additionally note that the decapping assay is performed in the absence of ATP. This means that CsdA helicase activity is not necessary for CsdA to modulate NudC cleavage of NAD^+^ caps.

**Figure 5.**
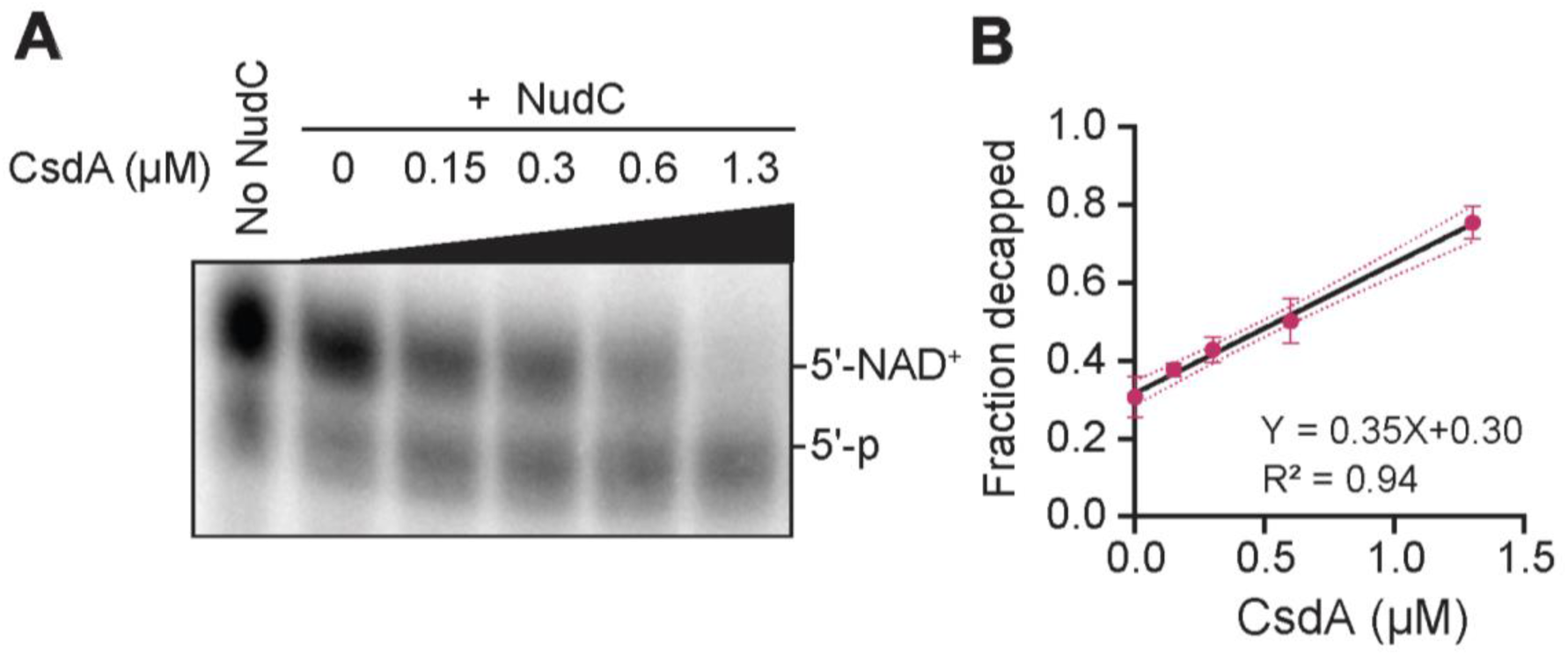
CsdA enhances the decapping efficiency of NudC. (**A**) Decapping activity of NudC on a single stranded NAD^+^ capped RNA in presence of CsdA *in vitro*. NAD^+^ capped probe RNA was incubated with 0.5 µM of NudC and increasing amounts of CsdA (as indicated) at 37°C for 30 mins. Top band indicates 5′ NAD^+^ capped RNA, and lower band indicates decapped RNA with a 5′ monophosphate end. (**B**) Quantification of NudC decapping shown in (A). Dots indicate fraction of decapped RNA for each CsdA concentration (as indicated). Data represent three independent replicates. Points indicate mean values, and error bars represent standard deviation. Statistical analysis was performed using simple linear regression to get the best-fit line. Line fit and R2 value is noted.

## Discussion

We have used *E. coli* NAD cap RNA affinity purification (NcRAP) to identify the first protein that interacts with a 5′ NAD^+^ capped RNA without enzymatically acting on it: the DEAD box RNA helicase, CsdA (**Fig. 1A**). In addition to CsdA, we also identified Hfq and ribosomal protein S1 as putative NAD^+^ cap RNA interacting protein. A recent study (19) showed that the only previously known NAD^+^ cap RNA interacting protein in *E. coli*, NudC, physically interacts with both CsdA and Hfq. Notably, we did not identify NudC in our pulldown. This result is surprising in context of NudC’s putative role as the primary NAD^+^ decapping enzyme through its NADH hydrolase activity (^5,35^). However, previous work by Höfer and colleagues noted that overexpressed wildtype and a catalytically inactive mutant (E178Q) NudC co-purifies with cellular RNA. They sequenced this RNA and found that it had a nearly identical composition to total cell RNA (^5^). This may indicate that NudC was not enriched in our NAD^+^ RNA pulldown assay because it non-specifically interacts with all RNA, independent of the presence of an NAD^+^ or another metabolite cap.

The identification of both the small regulatory RNA chaperone Hfq and ribosomal protein S1 as putative NAD cap interactors suggests an important role for NAD^+^ caps in post-transcriptional regulation of gene expression. This is consistent with findings that the primary transcripts in *E. coli* that carry NAD^+^ caps are small regulatory RNAs (1,4). Furthermore, ribosomal protein S1’s role in interacting with the leader sequence at 5′ UTR of mRNA, suggests that it may be able to differentiate between NAD^+^ capped and uncapped mRNAs during translation initiation (36). Both of these possibilities will require additional studies to better understand how these proteins facilitate downstream activities upon interacting with NAD^+^ capped RNAs. Our characterization demonstrates that CsdA is the first identified NAD^+^ cap “reader” protein in *E. coli*.

CsdA is an ATP dependent RNA helicase, which is well studied for its role in ribosome biogenesis at 37°C and its interaction with RNase E in low temperature conditions (15,18). Given these diverse functions, the ability of CsdA to interact with NAD^+^ capped RNAs is of particular interest. Consistent with NcRAP, we found CsdA has 8-fold higher affinity for NAD^+^ capped RNA over 5′ ppp RNA, along with differential binding kinetics (**Table 1**). A slower k_off_ value and higher binding response (nm) for NAD^+^ RNA suggest the presence of a distinct 5′ NAD^+^ cap binding site in the protein compared to uncapped RNAs. In line with this, we observed that CsdA preincubated with ppp RNA still binds NAD^+^ RNA, but less with ppp RNA binding when pre-incubated with NAD^+^ RNA (**Fig. 2E, F**). In addition, we found that a sub saturating concentration of NAD^+^ capped RNA stimulated higher ATPase activity of CsdA (**Fig. 3A-C**) relative to uncapped RNA. The ATPase function of CsdA is reported to be crucial for its RNA helicase activity (31,32). In line with a previous study, we observed that dsRNA substrates reduced the overall ATPase activity of CsdA (**Fig. 3B, C**), likely due to increased thermal stability of the RNA constructs (32). However, the relatively higher ATPase activity for double stranded NAD capped RNAs at sub-saturating concentrations suggests that preferential interaction with these capped substrates, allows more efficient initiation of helicase activity compared to uncapped RNA species. Collectively, these findings suggest that the presence of 5′ NAD^+^ cap, together with 5′ secondary structure and/or primary sequence, may contribute to selective recruitment of low-abundance NAD^+^ capped RNAs by CsdA for subsequent downstream activities.

Previous work has shown that *E. coli* CsdA can form condensates with RNA both *in vivo* and *in vitro* (33). Our findings indicate 5′ NAD^+^ cap might be an additional critical factor that helps in the formation of these condensates. The NAD^+^ cap likely introduces additional multivalent RNA-RNA or RNA-CsdA interactions, either directly through hydrophobic and π–π contacts of the nicotinamide ring with the protein, or indirectly through cap-induced changes in RNA structure. Invariably, the binding kinetics from BLI experiments (**Fig. 2C, D**) show that NAD^+^ cap confers a higher RNA-CsdA binding affinity that likely increases the effective population of RNA-CsdA complexes that can drive phase separation at a lower RNA concentration than is necessary with ppp RNA. From an *in vivo* perspective, this suggests that CsdA interaction with NAD^+^ capped RNAs could facilitate phase separation. Future studies will be necessary to understand if this 5′ NAD^+^ cap dependent condensate formation by CsdA allows specific transcripts or proteins to be recruited at the site for further downstream activities.

CsdA has been reported to physically interact with NudC (19). Consistent with this finding, we found that CsdA facilitates NAD^+^ cap cleavage by NudC (**Fig. 5**). CsdA increases this decapping activity of NudC independent of CsdA’s ATPase activity. This suggests that this function of CsdA is independent of its RNA unwinding activity, which requires ATP hydrolysis. Surprisingly, this also means that the strong binding between NAD^+^ capped RNA and CsdA does not block accessibility of 5′ NAD^+^ cap to NudC. However, we do not know how the RNA condensate forming of activity may affect CsdA’s interaction with NudC or the stimulation of NAD^+^ decapping. It is possible that NudC is present in these condensates, but it may also be that condensate formation may sequester NAD^+^ capped RNAs, protecting them from NudC activity. Further experiments will be required to determine if NudC is present and enzymatically active in these condensates.

Decapping by NudC generates 5′ monophosphate ends and has been proposed to be the first step in degradation of capped transcripts (^5^). This 5′ monophosphate can be recognized and further degraded by RNase E in the RNA degradosome. A previous study has reported that CsdA interacts with RNase E at low temperatures (18). Taken together with our characterization of CsdA, this suggests that CsdA likely plays an important role in regulating degradation of NAD^+^ capped transcripts. Further investigation is required to determine if these interactions also occur under normal growth temperature (37°C). Overall, our results indicate that the preferential interaction between CsdA and NAD⁺-capped RNAs plays a key role in maintaining NAD⁺ RNA homeostasis in *E. coli*.

## AUTHOR CONTRIBUTION

Conceptualization, K.D, J.M.S, W.S.C. and J.G.B.; Methodology, K.D., K.G.D., Y.S., Y.Y., K.R.S., J.M.S., W.S.C., and J.G.B.; Formal Analysis, K.D., Y.Y., K.S., and J.G.B.; Investigation, K.D., K.G.D. and Y.S.; Writing, K.D., K.R.S and J.G.B.; Visualization, K.D. and J.G.B.; Supervision, W.S.C., and J.G.B.; Project Administration, J.G.B.; Funding Acquisition, J.G.B.

## SUPPLEMENTARY DATA

Supplementary Data are available at NAR online.

## CONFLICT OF INTEREST

The authors declare no conflict of interest.

## FUNDING

This work was supported by the University of Delaware Research Foundation (24A01689), and the NIH National Institute of General Medical Sciences (P20GM104316).

## DATA AVAILABILITY

All data underlying this study are provided in the article and its online supplementary material. The MS raw files associated with this study have been deposited to the MassIVE server (https://massive.ucsd.edu/) with the dataset identifier MSV000102001 (and passcode a).

## SUPPLIMENTAL MATERIALS

**Supplementary Figure 1.**
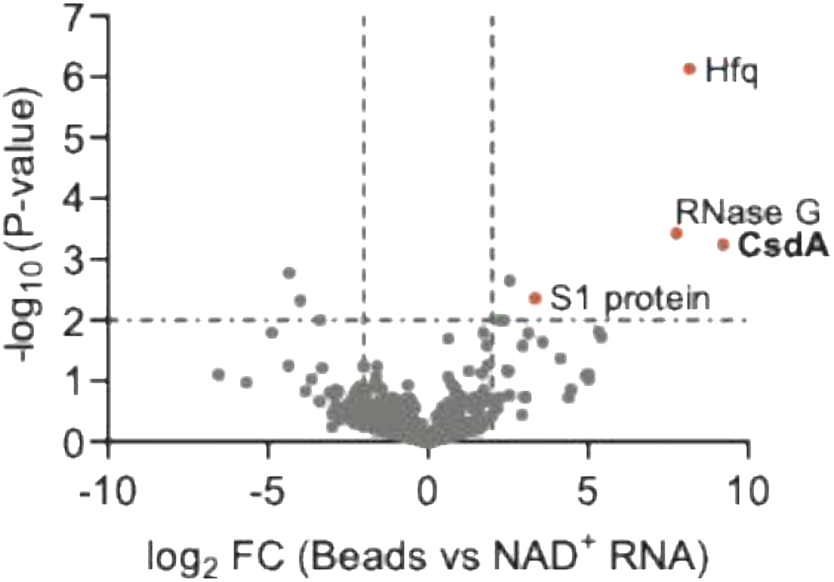
Volcano plot of proteins differentially enriched in NAD^+^ capped RNA vs Beads alone pulldowns from NcRAP. The x-axis corresponds to log_2_ fold change of protein association (NAD^+^ RNA vs Beads); while the y-axis shows -log_10_ transformed P values. Dashed lines indicate significance threshold defined by log_2_ fold change > 2 and -log _10_ P value > 2; proteins meeting both criteria were considered significantly enriched. Statistical analysis was performed using two-sided t-test for comparison.

**Supplementary Figure 2.**
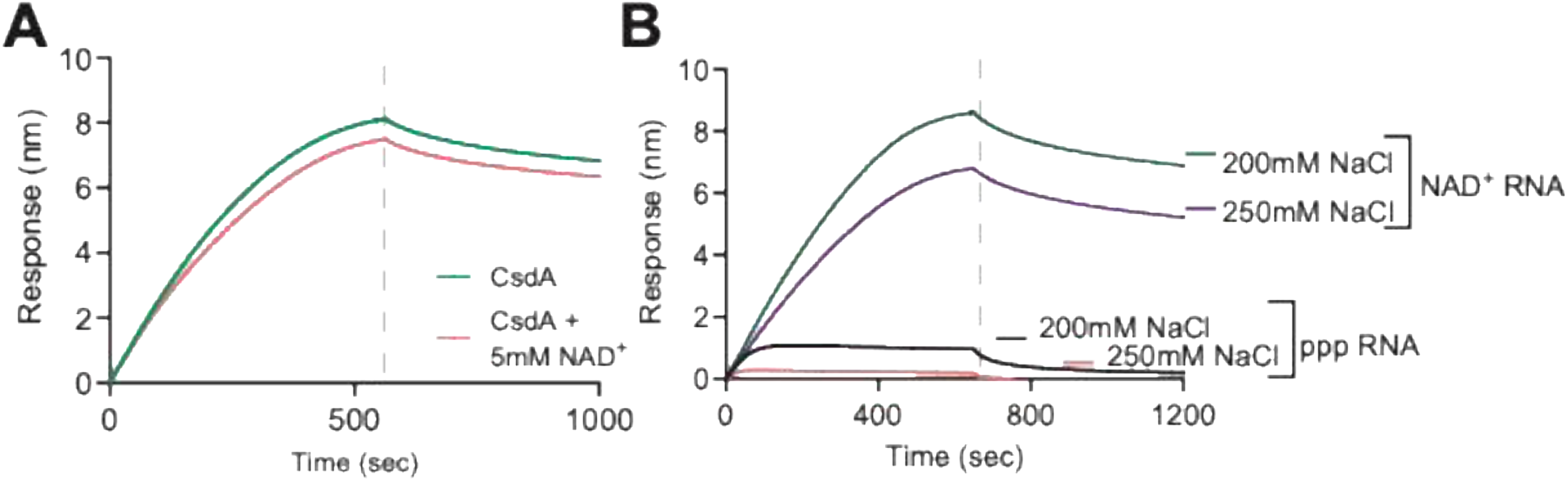
Influence of NAD^+^ and salt concentration on CsdA interaction with NAD^+^ capped RNA using Biolayer interferometry. (**A**) Biosensor response (nm) when purified CsdA interacts with NAD^+^ capped RNA in presence and absence of 5mM NAD^+^. (**B**) Biosensor response when 50 nM CsdA interacts with 50 nM NAD^+^ or ppp RNA in presence of varying salt concentrations.

**Supplementary Figure 3.**
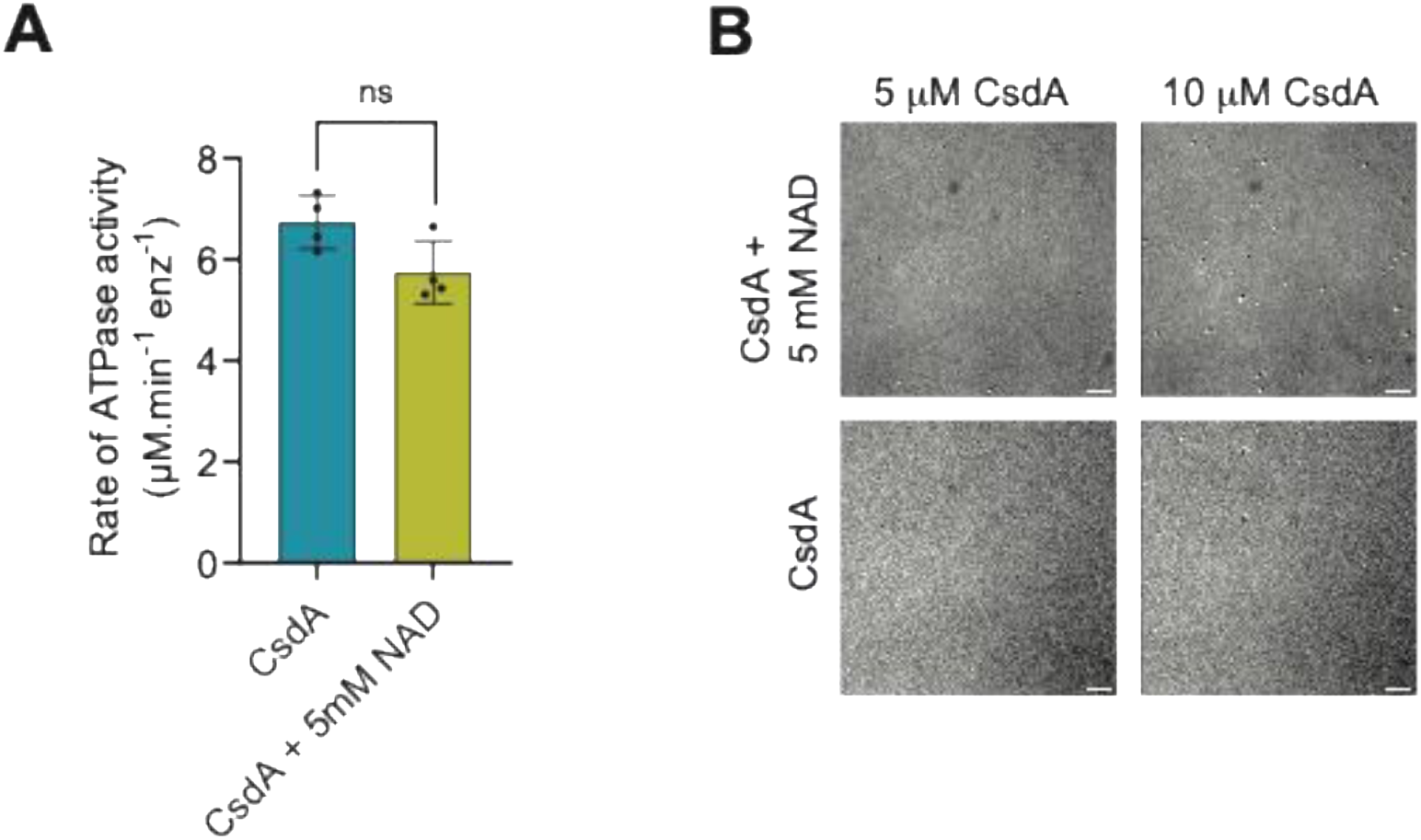
Effect of NAD^+^ on CsdA’s ATPase and condensate formation activity. (**A**) ATPase activity of CsdA measured in the absence of RNA and in presence of 5mM NAD^+^ (**B**) Phase separation of purified CsdA in absence of RNA and in presence of 5 mM NAD^+^.

**Supplementary Figure 4.**
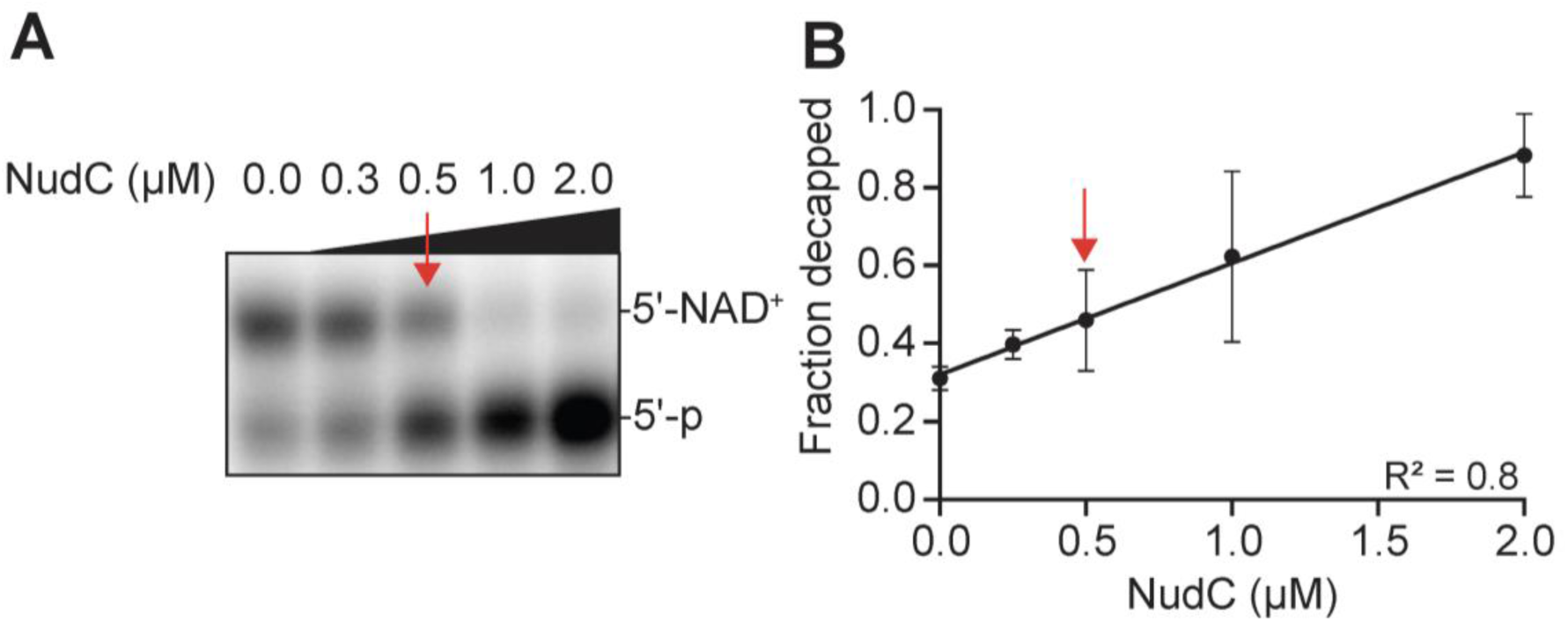
Decapping activity of NudC with NAD^+^ capped RNA (**A**) Decapping of *in vitro* synthesized NAD^+^ probe RNA when incubated with varying amounts of NudC (as indicated) at 37°C for 30 mins. (**B**) Quantification of the gel image in A from three independent experiments. Points indicate mean values. Error bars denote standard deviation. Arrow indicates 0.5 µM NudC concentration, which was used for Fig 5A (see main text). Statistical analysis was performed using simple linear regression to get the best fit line.

## MATERIALS AND METHODS

**Supplemental table 1.** Reagents.

| Items | Source | Catalog number |
| --- | --- | --- |
| Nuclease-Free water | Thermo Fisher Scientific | AM9932 |
| NAD<sup>+</sup> | Sigma Aldrich | 10127965001 |
| NADH | Sigma Aldrich | 10128023001 |
| Phosphoenol-pyruvate | Sigma Aldrich | 10108294001 |
| ATP | GoldBio | A-081-1 |
| Biocytin | Sigma Aldrich | B4261-5MG |
| Anhydrotetracycline | Fisher Scientific | AC233111000 |
| Isopropyl β-D-1-thiogalactopyranoside (IPTG) | GoldBio | I2481C25 |
| TRI Reagent | Molecular Research Center | TR118 |
| SYBR Gold nucleic acid gel stain | Thermo Fisher Scientific | S11494 |
| SimplyBlue SafeStain | Thermo Fisher Scientific | LC6060 |
| NEBExpress *E. coli* lysis Reagent | New England Biolabs | P8116S |
| Ni-NTA Agarose | MCLAB | NINTA-25 |
| Amicon Ultra Centrifugal Filter, 30kDa MWCO | Sigma Aldrich | UFC903024 |
| Dynabeads Myone C1 Streptavidin Magnetic Beads | Thermo Fisher Scientific | 65001 |
| TURBO DNase | Thermo Fisher Scientific | AM2238 |
| T4 DNA ligase | New England Biolabs | M0202S |
| T4 RNA ligase 1<br>(ssRNA ligase) | New England Biolabs | M0204S |
| NEBExpress T4 lysozyme | New England Biolabs | P8115S |
| Pyruvate Kinase/Lactic<br>Dehydrogenase | Sigma Aldrich | P0294-5ML |
| Anti-FLAG M2 antibody mouse<br>monoclonal | Sigma Aldrich | F3165-2MG |
| Goat anti-Mouse IgG<br>Secondary antibody, HRP | Thermo Fisher Scientific | 31431 |
| pCP-Biotin | Jena Bioscience | NU-1706-BIO |
| HiScribe T7 High yield RNA<br>Synthesis Kit | New England Biolabs | E2040S |
| Direct-zol RNA Miniprep plus kit | Zymo Research | R2072 |
| Phenol: Chloroform:Isoamyl<br>alcohol (25:24:1) | Sigma Aldrich | P3803-400ML |
| Novex TBE Urea gel, 10% | Thermo Fisher Scientific | EC6875BOX |
| NuPAGE Bis-Tris Mini protein<br>gels, 4%-12% | Thermo Fisher Scientific | NP0321BOX |
| TransBlot Turbo Mini-size<br>nitrocellulose 7.1*8.5 cm | Bio-Rad | 1704270 |
| Clarity Western ECL substrate | Bio-Rad | 1705061 |
| NuPAGE LDS sample buffer | Thermo Fisher Scientific | NP0007 |
| α-32P GTP 0.25mCi | Perkin Elmer | BLU506H250UC |
| Bioruptor Plus | Diagenode | B01020014 |
| E3 filters | Fisher Scientific | 13-110-084 |
| C18 stage tips | Fisher Scientific | 13-110-056 |
| Immersol Immersion oil 518F | Fisher Scientific | NC1997771 |
| Octet Streptavidin (SA)<br>Biosensor | Sartorius | 18-5019 |
| Nikon TiE inverted microscope | Nikon | - |
| Andor Ixon Ultra 897 EMCCD camera | Oxford Instruments | - |
| AKTA Pure | Cytiva | - |
| SPARK multimode microplate reader | Tecan Life Sciences | - |
| Octet RH16 system | Sartorius | - |
| Orbitrap Eclipse Tribrid Mass spectrometer | Thermo Fisher Scientific | - |

**Supplemental Table 2.** Biological Resources.

| **Strains** | **Genotype/Phenotype** | **Source** |
| --- | --- | --- |
| *Escherichia coli* | MG1655 | ATCC (#47076) |
| *Escherichia coli* | MG1655 *csdA*::3x FLAG | This study |
| *Escherichia coli* | NiCo 21 (DE3) | New England Biolabs (#C2529H) |
| **Plasmid** | **Construct** | **source** |
| pET28c-CsdA-His | DNA fragment encoding full length *csdA*, flanked by HindIII and Xba1 sites, was cloned into pET28c to generate a C-terminal polyhistidine (6x HIS) tagged construct | This study |
| pET28C-NudC-His | DNA fragment encoding full length *nudC*, flanked by HindIII and Xba1 sites, was cloned into pET28c to generate a C-terminal polyhistidine (6x His) tagged construct. | Bird, J. G et al. (2016) *Nature* |
| pET28C-NudCE178Q -His | DNA fragment encoding full length *nudC* with an amino acid mutation E178Q, flanked by HindIII and Xba1 sites, was cloned into pET28c to generate a C- | This study |
|  | terminal polyhistidine (6x His) tagged construct. |  |
| pSLTS | DNA fragment for expression of $\lambda$ red recombinase and *Isce1* endonuclease | Addgene (#59386) |
| pCP20 | DNA fragment for the expression of FLP recombinase | Gift from Dr Bryce Nickels (Rutgers University) |

**Supplemental Table 3.**
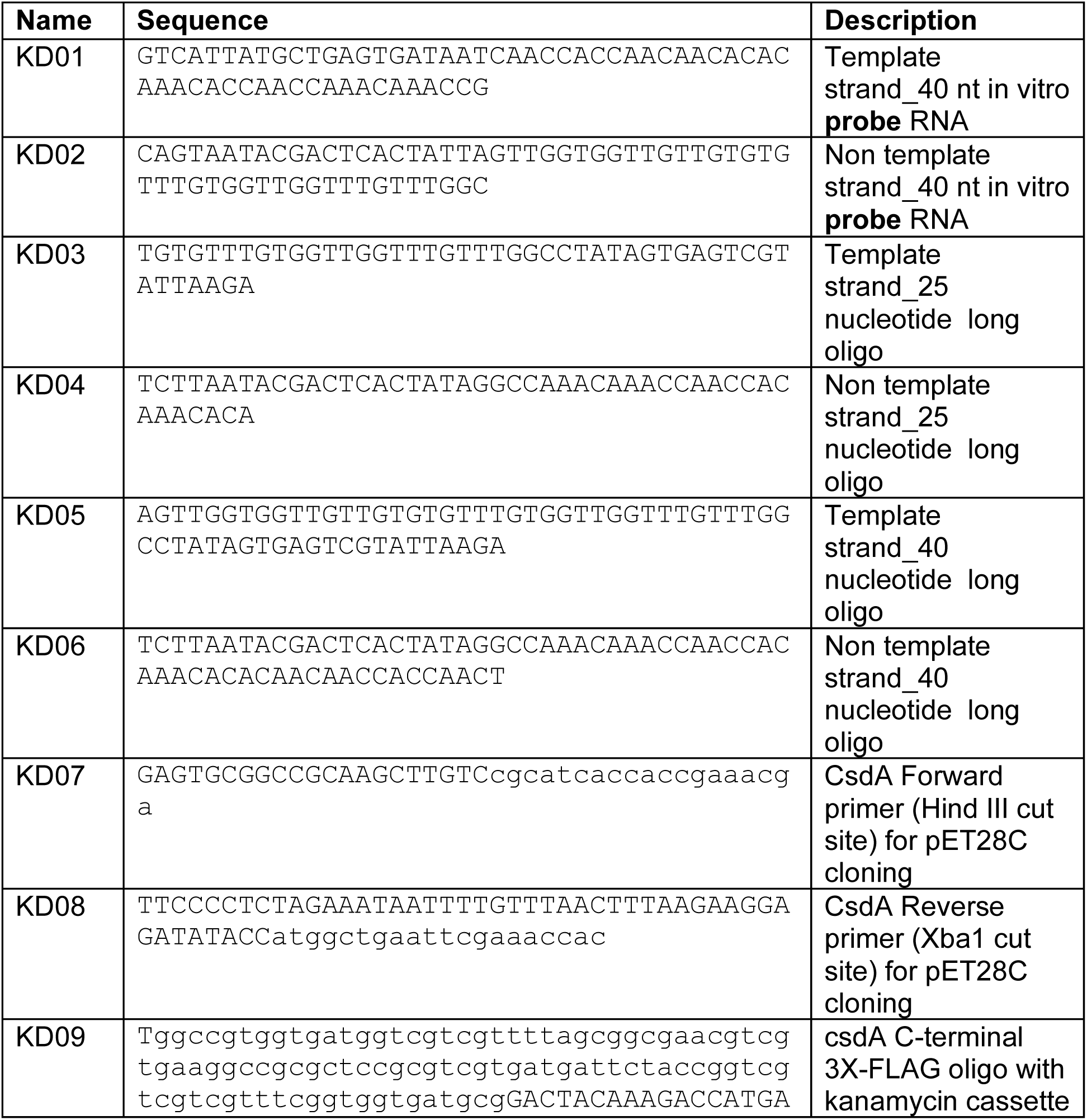
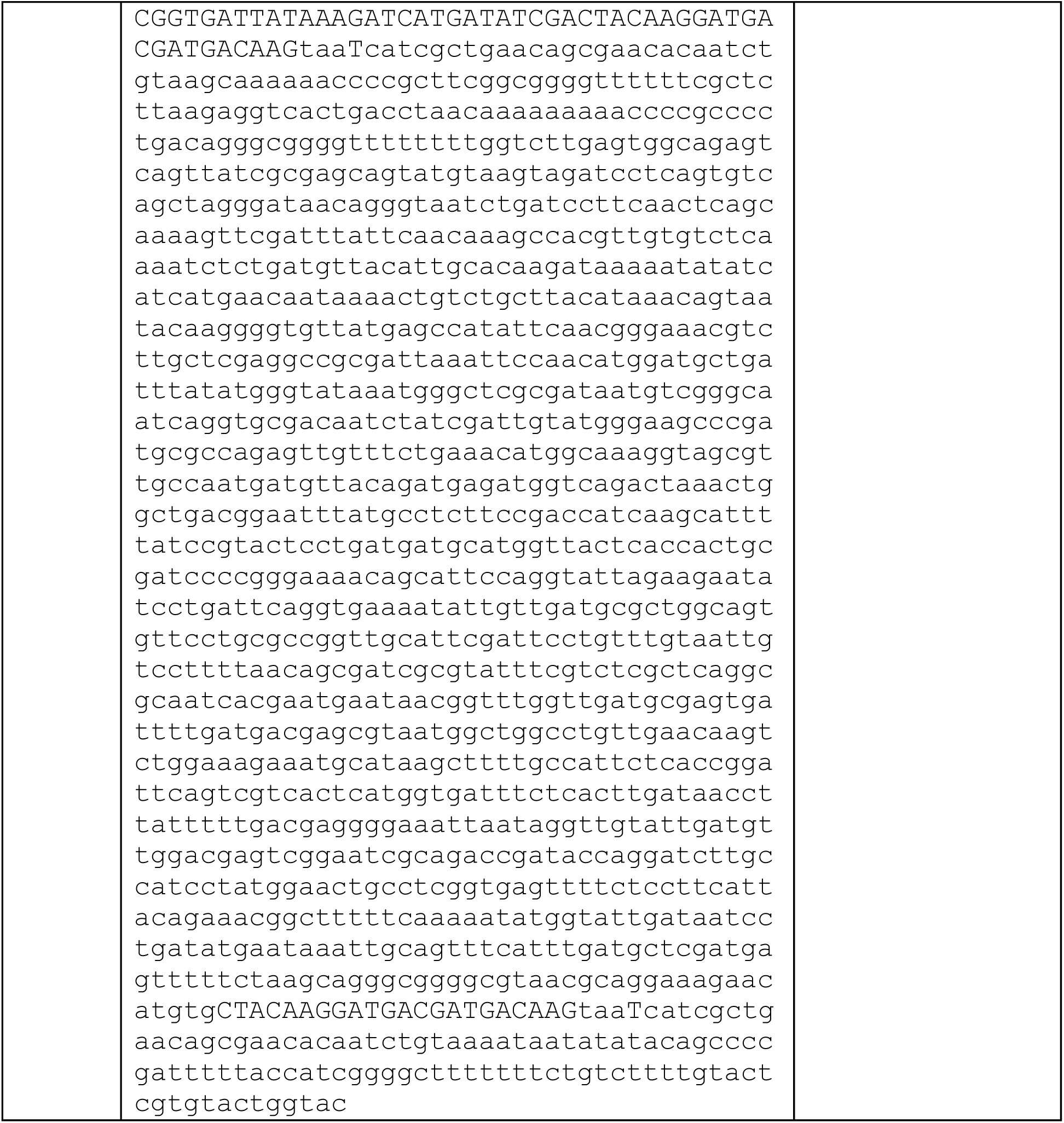
Synthesized Oligonucleotides used in this work.

## Notes

### Competing Interest Statement

The authors have declared no competing interest.

